# Crystal structures and functional analysis of the ZnF5-WWE1-WWE2 region of PARP13/ZAP define a new mode of engaging poly(ADP-ribose)

**DOI:** 10.1101/2021.12.15.472832

**Authors:** Jijin R. A. Kuttiyatveetil, Heddy Soufari, Morgan Dasovich, Isabel R. Uribe, Shang-Jung Cheng, Anthony K. L. Leung, John M. Pascal

## Abstract

PARP13/ZAP acts against multiple viruses through recognizing and promoting degradation of cytoplasmic viral mRNA. PARP13 has four N-terminal Zn-finger motifs that bind CG-rich nucleotide sequences, and a C-terminal ADP ribosyltransferase fold similar to other PARPs. A central region predicted to contain a fifth Zn-finger and two tandem WWE domains is implicated in binding poly(ADP-ribose); however, there are limited insights into the structure and function of this PARP13 region (ZnF5-WWE1-WWE2). Here, we present crystal structures of ZnF5-WWE1-WWE2 from mouse PARP13 in complex with ADP-ribose and with ATP. ZnF5-WWE1-WWE2 crystallized as a dimer with major contacts formed between WWE1 and WWE2 originating from different monomers, indicative of a more compact monomeric arrangement of the tandem WWE domains. Solution scattering experiments and biophysical analysis indicated a monomer in solution, suggesting that the crystal dimer represents domain swapping that could potentially represent a PARP13 conformation assumed when signaling viral RNA detection. The crystal structure and binding studies demonstrate that WWE2 interacts with ADP-ribose and ATP, whereas WWE1 does not have a functional binding site. The shape of the WWE2 binding pocket disfavors interaction with the ribose-ribose linkage of poly(ADP-ribose). Binding studies with poly(ADP-ribose) ligands indicate that WWE2 serves as an anchor for preferential binding to the terminal end of poly(ADP-ribose), and the composite structure of ZnF5-WWE1-WWE2 forms an extended surface to engage polymer chains of ADP-ribose. This model represents a novel mode of poly(ADP-ribose) recognition and provides a structural framework for investigating poly(ADP-ribose) impact on PARP13 function.

## INTRODUCTION

PARP13 belongs to the PARP protein family that has in common an ADP ribosyl transferase (ART) fold that uses NAD^+^ to modify proteins and nucleic acids with ADP-ribose (Hottiger *et al*., 2010; Lüscher *et al*., 2018; Vyas *et al*., 2014). The product of PARP reactions is primarily modulated through ART active site variations that result in certain family members producing poly(ADP-ribose) modifications, most family members producing mono-ADP-ribose modifications, and some members lacking ADP-ribosyl transferase activity entirely, such as PARP13 (Kleine *et al*., 2008; Hottiger *et al*., 2010; Vyas *et al*., 2014; Karlberg *et al*., 2015; Lüscher *et al*., 2018). Poly(ADP-ribose) and ADP-ribose modifications act in a variety of manners to coordinate cellular processes, such as the DNA damage response, gene regulation, cellular signaling, and the antiviral response.

PARP13 was first known as zinc-finger antiviral protein (ZAP) due to the presence of zinc-fingers and activity against retroviral RNA (Gao, Guo and Goff, 2002). The antiviral response is accomplished through PARP13 recognition of cytoplasmic mRNA, which recruits RNA degradation machinery through multiple mechanisms to eliminate target viral mRNA (Guo *et al*., 2007; Zhu and Gao, 2008). PARP13 overexpression is associated with replication inhibition of viruses such as Filovirus, Alphaviruses and numerous others (Bick *et al*., 2003; Müller *et al*., 2007; Mao *et al*., 2013; Liu *et al*., 2015). Human PARP13/ZAP has four splice variants of varying lengths, with the major difference being that the two shortest isoforms lack the ART domain entirely (Li *et al*., 2019). While all four isoforms exhibit similar antiviral activities against most virus groups, the longer isoforms containing the ART fold exhibit greater antiviral potential against alphaviruses and hepatitis B virus (HBV).

The N-terminal domain of PARP13 has four CCCH-type Zn-finger motifs that collectively bind CG-rich RNA sequences (Gao, Guo and Goff, 2002; Guo *et al*., 2004; Chen *et al*., 2012). Recently, crystal structures of human and mouse PARP13 N-terminal zinc-fingers bound to RNA have been determined, revealing the structural basis of RNA recognition (Meagher *et al*., 2019; Luo *et al*., 2020). The structures clearly signify flexibility in the regions of the Zn-fingers that interact with single-strand RNA, and the presence of a highly electropositive RNA binding path that recognizes CG dinucleotides. Interestingly, the CG dinucleotide binding pocket of the second Zn-finger is highly specific and discriminates other nucleotides, while the rest of the contacts with single-strand RNA are not specific. It is postulated that multiple PARP13 molecules will bind to regions of viral RNA rich in CG dinucleotides, and the assembly of multiple PARP13 will then act as a signal to recruit the mRNA destruction complex (Luo *et al*., 2020). However, the molecular mechanism for PARP13 signaling of RNA recognition is not understood and will likely require further insights into the structure of PARP13.

Compared to the N-terminal zinc-fingers and the C-terminal ART, much less is known about the structure and function of the central region of PARP13, which is present in all four isoforms. This central region contains ~270-residues of low complexity sequence that are likely unfolded, followed by ~220-residues just before the ART domain, that are predicted by sequence to contain a zinc finger and one or two WWE domains. WWE domains (named for conserved residues Trp-Trp-Glu) are protein modules known to form protein-protein interactions or to bind to poly(ADP-ribose) (Wang *et al*., 2012). Indeed, PARP13 has been identified as a poly(ADP-ribose)-binding protein in proteomic analyses (Dasovich *et al*., 2021; Kliza *et al*., 2021). We undertook a structural and functional analysis of the central PARP13 region predicted to contain folded structures, as none of these features had been confirmed. It was also unclear whether the predicted domains formed a single globular unit or a collection of protein modules. Moreover, the properties of the potentially tandem WWE domains was intriguing, as the best studied WWE domain that is known to interact with poly(ADP-ribose), RNF-146, utilizes a single WWE domain to bind the ribose-ribose linkage of poly(ADP-ribose) (Wang *et al*., 2012). Our crystallographic and solution-based biophysical analysis collectively indicates a compact, monomeric fold for the central region of PARP3, but also hints at a level of structural plasticity in the zinc finger fold and a potential to form higher order assemblies. The crystal structures highlight unique ADP-ribose interaction features in a binding pocket in the second WWE domain of PARP13. Together with binding studies, we present a novel mode of engaging poly(ADP-ribose) from an anchor point on the terminal ADP-ribose unit.

## RESULTS

### Production of ZnF5-WWE1-WWE2

Several human PARP13 (hP13) domain boundaries were tested for soluble protein production, including multiple constructs of ZnF5-WWE1-WWE2, constructs for individual domains (ZnF5, WWE1, or WWE2), and constructs for tandem WWEs (WWE1-WWE2). Three constructs for ZnF5-WWE1-WWE2 (residues 468-699, 487-699 and 507-699) produced soluble protein, and the construct coding for residues 507 to 699 was selected to represent human PARP13 (hP13) ZnF5-WWE1-WWE2, as it included all predicted domains in a minimal sequence (Figure 1A). None of the constructs designed to produce individual domains or domain combinations produced soluble protein that would permit further studies. Together, the expression studies indicated that the domains were best produced in combination, suggesting that they form a globular assembly. Zinc content analysis was performed on purified hP13 ZnF5-WWE1-WWE2 to verify the ZnF5 fold (Figure 1B). The results indicated that ~97% of the protein co-purified with zinc, consistent with the prediction of a fifth zinc finger domain in hP13. RNF-146 does not contain a zinc finger and served as a negative control, and the third zinc finger of PARP1 served as a positive control, both verifying the zinc content analysis procedure (Figure 1B).

**Figure 1:**
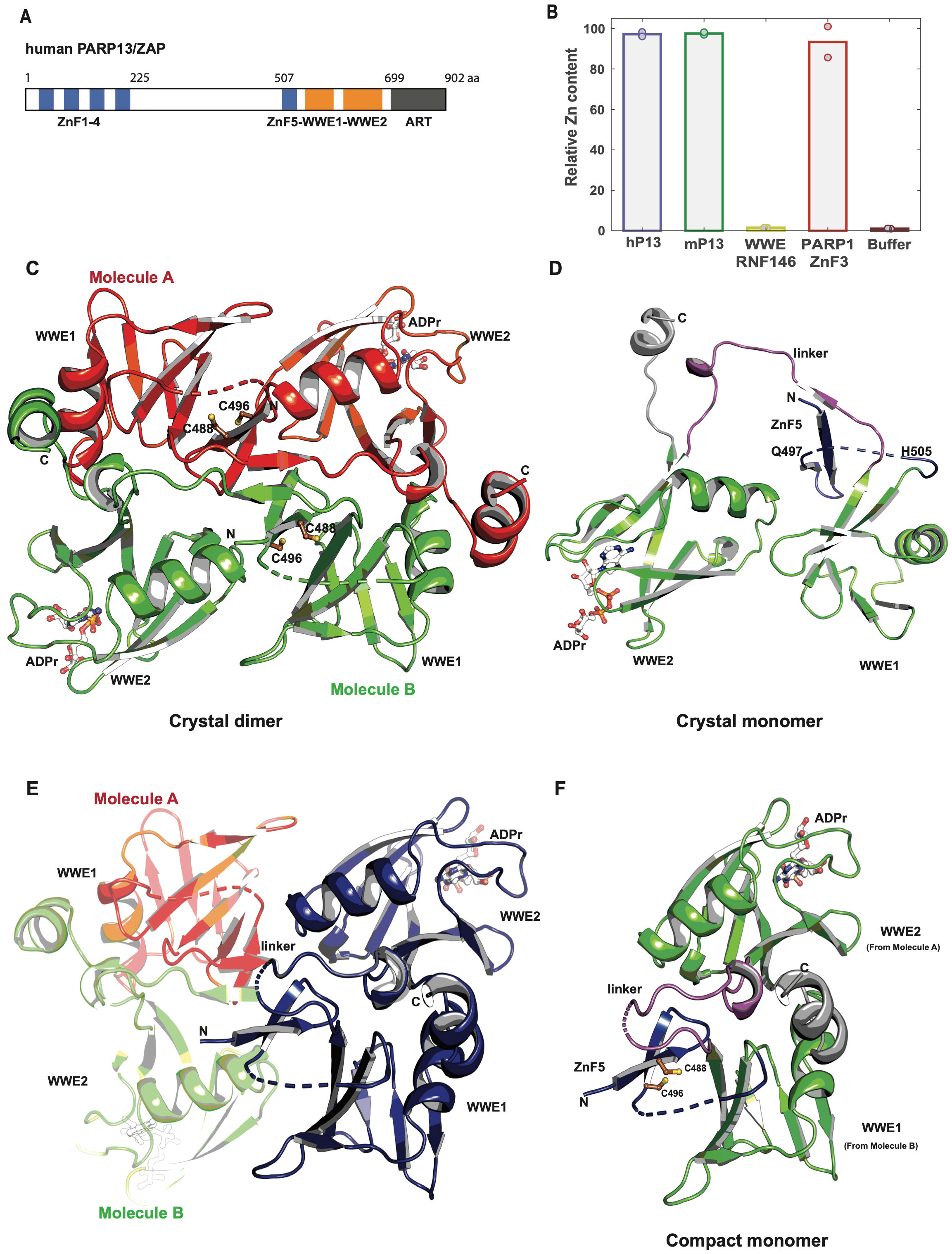
Crystal structure of mP13 ZnF5-WWE1-WWE2 bound to ADPr. (A) Domain organization of human PARP13/ZAP. (B) Zinc content analysis was performed on the designated samples, and the relative concentrations between samples are shown for two measurements each. Proteins were analyzed at 10 μM. A zinc concentration of 10 μM was set as 100, such that the values for protein samples represent the percent of molecules with bound zinc. RNF-146 is a control protein that does not contain a zinc finger; ZnF3 of human PARP1 is a control protein that does contain a zinc finger; and the dialysis buffer was used to estimate background levels of zinc. (C) Crystal dimer. The crystallographic asymmetric unit contains two molecules of mP13 ZnF5-WWE1-WWE2 (red Molecule A, and green Molecule B). Both molecules have an ADPr molecule bound to WWE2. Residues C488 and C496 form part of ZnF5, which is partially ordered in this structure. (D) Crystal monomer. One ZnF5-WWE1-WWE2 molecule from the asymmetric unit is shown, with the extended linker (purple), WWE domains (green), ZnF5 (deep blue), and residues leading to the CAT domain (grey) labeled. (E) Compact monomer. Crossing over between the linker regions of the two protein molecules (small dashed-line labeled “linker”) models a compact arrangement of WWE1 and WWE2 from different chains (compact monomer in blue). (F) Model for the compact monomer with the modified linker in purple, the WWE domains in green, ZnF5 region in deep blue, and residues leading to the CAT domain in grey.

Numerous crystallization experiments conducted with the different hP13 ZnF5-WWE1-WWE2 constructs in the presence and absence of ligands failed to yield crystals. We also cloned and produced the same ZnF5-WWE1-WWE2 region from mouse PARP13 (mP13; residues 476 to 673), which shares 60% identity with hP13 over this region. Zinc content analysis of mP13 also verified co-purification of stoichiometric amounts of zinc (Figure 1B), consistent with a zinc finger structure. mP13 ZnF5-WWE1-WWE2 yielded diffraction quality crystals in the presence of ATP and in the presence of ADPr. The crystal structures were determined by single-wavelength anomalous diffraction using seleno-methionine containing protein, with resolutions of 2.2 Å for the ADPr complex and 2.2 Å for the ATP complex (Table 1; Crystallographic statistics).

**Table 1.** Crystallographic data and refinement statistics.

| **Data Collection<sup>a</sup>** |  |  |
| --- | --- | --- |
| Structure | **mP13 ZnF5-WWE1-WWE2: ATP** | **mP13 ZnF5-WWE1-WWE2: ADPr** |
| PDB ID | 7SZ2 | 7SZ3 |
| Space Group | P3<sub>1</sub>21 | P3<sub>1</sub>21 |
| Unit Cell Dimensions | $a=b=87.54 \text{ \AA}$, $c=129.83 \text{ \AA}$<br>$\alpha=\beta=90^\circ$ $\gamma=120^\circ$ | $a=b=87.42 \text{ \AA}$, $c=128.74 \text{ \AA}$<br>$\alpha=\beta=90^\circ$ $\gamma=120^\circ$ |
| Wavelength (Å) | 1.1806 | 1.1806 |
| Resolution range (Å) | 43.77–2.20 (2.27–2.20) | 49.04–2.20 (2.27–2.20) |
| Completeness (%) | 100.0 (100.0) | 100.0 (99.9) |
| Unique observations | 29,855 (2,545) | 29,530 (2,519) |
| Average Redundancy | 21.8 (22.5) | 21.9 (22.4) |
| Mean (I/σI)<sup>b</sup> | 33.8 (2.3) | 28.7 (2.0) |
| R<sub>merge</sub> (%)<sup>b</sup> | 6.2 (163.9) | 8.5 (227.0) |
| R<sub>pim</sub> (%)<sup>b</sup> | 1.4 (35.2) | 1.8 (48.3) |
| Mean I CC(1/2)<sup>b</sup> | 1.0 (0.828) | 1.0 (0.849) |
| **Model Refinement<sup>a</sup>** |  |  |
| Resolution Range (Å) | 20–2.20 (2.25–2.20) | 20–2.20 (2.25–2.20) |
| Number of reflections | 27,170 (1958) | 26,802 (1915) |
| R<sup>c</sup> | 0.2027 (0.319) | 0.1926 (0.380) |
| R<sub>free</sub><sup>c</sup> | 0.2384 (0.340) | 0.2305 (0.367) |
| Number of atoms /<br>Average B-factor (Å<sup>2</sup>) | 3,236 / 66.0 | 3,230 / 63.0 |
| Protein | 3,003 / 65.9 | 2,998 / 62.6 |
| Solvent | 171 / 75.5 | 160 / 72.20 |
| Ligand | 62 / 81.2 | 72 / 77.7 |
| Phi/Psi, preferred (%) /<br>outliers (#)<sup>d</sup> | 94.22 / 0 | 95.07 / 0 |
| R.m.s.d. bond angles (°) | 1.4375 | 1.5275 |
| R.m.s.d. bond lengths (Å) | 0.0068 | 0.0082 |
| <sup>a</sup> Values in parentheses refer to data in the highest resolution shell.<br><sup>b</sup> As calculated in SCALA: $R_{\text{merge}} = \sum_{hkl} \sum_j \vert I_j - \langle I \rangle\vert / \sum_{hkl} \sum_j I_j$. $\langle I \rangle$ is the mean intensity of $j$ observations of reflection $hkl$ and its symmetry equivalents; $R_{\text{pim}}$ takes into account measurement redundancy when calculating $R_{\text{merge}}$; Mean I CC(1/2) is the correlation between mean intensities calculated for two randomly chosen half-sets of the data.<br><sup>c</sup> $R = \sum_{hkl} \vert F_{\text{obs}} - kF_{\text{calc}}\vert / \sum_{hkl} \vert F_{\text{obs}}\vert $ for reflections used in refinement. $R_{\text{free}} = R$ for 5% of reflections excluded from crystallographic refinement.<br><sup>d</sup> As reported in COOT. | | |

### Overview of the mP13 ZnF5-WWE1-WWE2 crystal structure

mP13 ZnF5-WWE1-WWE2 crystallized with two essentially identical molecules in the asymmetric unit (Figure 1C, red and green molecules). The two molecules of mP13 ZnF5-WWE1-WWE2 form a two-fold symmetric dimer (Figure 1C), with an extensive buried surface area of over 5,000 Å^2^. Each molecule indeed contains two WWE folds, although with key differences distinguishing WWE1 and WWE2. Most notably, only WWE2 showed evidence of bound ADPr/ATP and WWE1 lacked a binding pocket. The ZnF5 residues anticipated to form a CCCH-type zinc finger (zinc-coordinating residues: C488, C496, C501, and H505) were only partially modeled due to a lack of electron density for certain regions, presumably due to disorder. The modeled region of ZnF5 is positioned adjacent to WWE1. In each of the two molecules, ZnF5-WWE1 is linked to WWE2 by a 15-residue, extended linker (one monomer is shown in Figure 1D). Following WWE2 is a small helix that would in turn lead to the ART domain in the longer isoforms of PARP13. Thus, each of the two molecules alone forms a fairly extended structure (Figure 1D).

### Analysis of WWE domain interactions

We noted that WWE1 and WWE2 domains from *different* molecules form a large surface area burying over 2000 Å^2^ (e.g., green WWE1, red WWE2 combination in Figure 1C), whereas WWE domains from the same chain form a smaller contact area less than 800 Å^2^ (see Figure 1D). We also noted that a simple crossing over between polypeptides in the extended linker near the axis of two-fold symmetry allowed us to model the more extensive WWE1/WWE2 interactions as arising from the same polypeptide (see crossing over in Figure 1E and resulting model in Figure 1F). The resulting configuration of the two WWE domains is substantially more compact than the extended configuration (Figure 1F versus Figure 1D, respectively). We viewed the compact configuration as more plausible based on the problems producing the WWE domains separately, which implied a more compact assembly of these two WWE domains. We pursued solution-based biophysical techniques to further examine ZnF5-WWE1-WWE2 structure, and as described in the next sections, the biophysical analysis strongly supports the compact monomer configuration. Moreover, the analysis addresses the conformation of the ZnF5 region that is only partially structured in our crystallized complexes.

### Biophysical analysis of ZnF5-WWE1-WWE2 in solution

We first analyzed the multimeric state of ZnF5-WWE1-WWE2 in solution. The molar mass of the ZnF5-WWE1-WWE2 region of mP13 and hP13 was evaluated using SEC-MALS (size exclusion chromatography/multi-angle light scattering) (Figure 2A). The elution profile for mP13 had two peaks: a major peak at ~16.5 mL representing 96% of the scattering mass and estimated at 25.7 kDa ± 0.5%, and a minor peak at ~15.3 mL representing ~4% of the scattering mass and estimated at 49 kDa ± 1.7%. These mass values best estimate a monomer (96%) and dimer (~4%) of mP13 ZnF5-WWE1-WWE2 (theoretical mass of 23.6 kDa). The elution profile for hP13 also exhibited two peaks. The larger peak at ~16.7 mL represented 98% of the scattering mass and was estimated at 25.1 kDa ± 0.4%. The scattering signal from the smaller hP13 peak was not sufficient to provide a mass estimate, but its location appears similar to the dimer peak observed in the mP13 sample. Therefore, both mP13 and hP13 exist primarily as monomers in solution, with some evidence for a small population of dimers.

**Figure 2:**
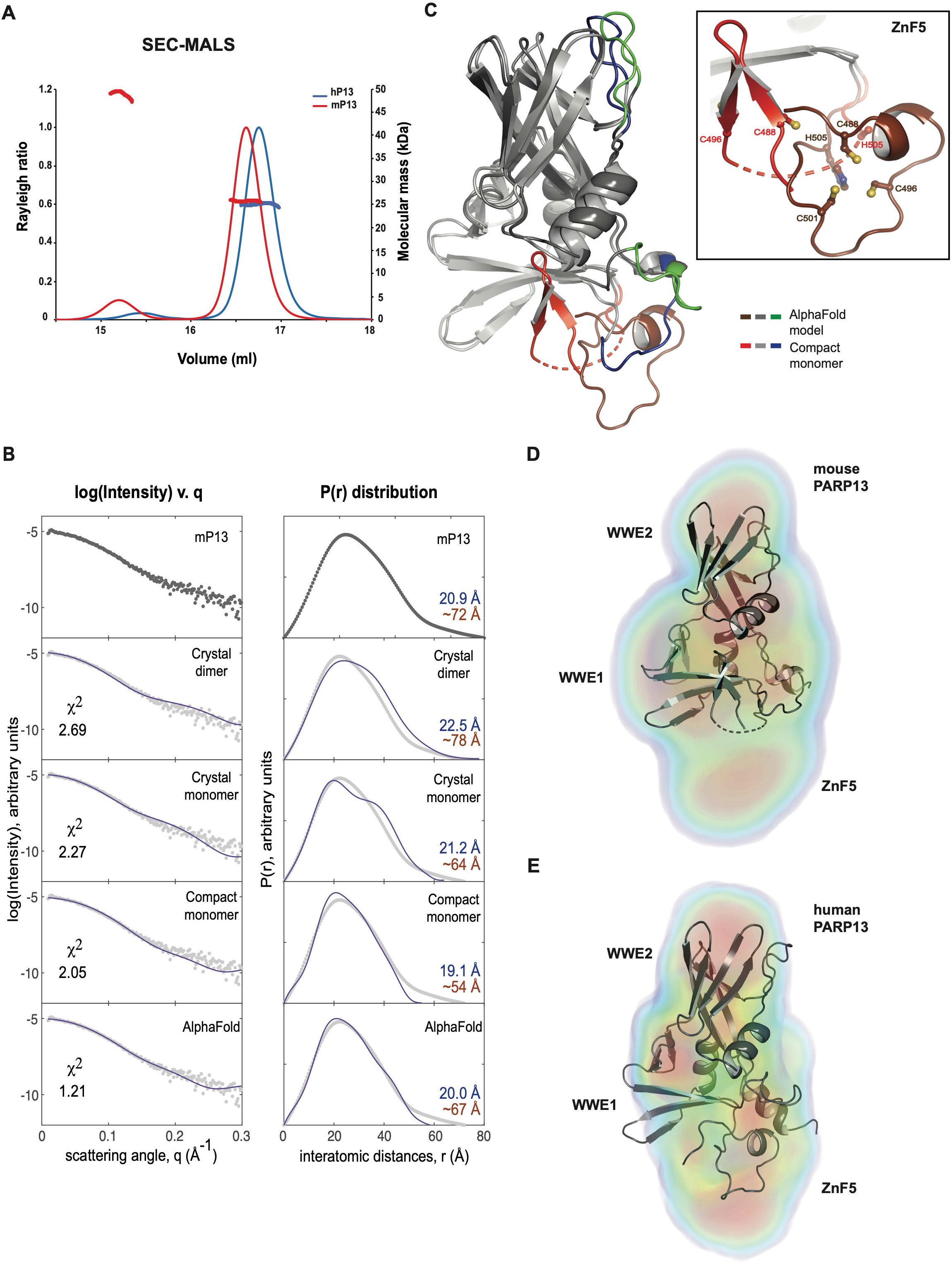
Biophysical analysis of PARP13 ZnF5-WWE1-WWE2. (A) SEC-MALS analysis of ZnF5-WWE1-WWE2 from hP13 (blue) and mP13 (red). The light scattering data is plotted as the Rayleigh ratio (left y axis) versus elution volume from gel filtration. The mass estimation (right y axis) is shown across the indicated peaks. (B) Left column. The experimental SAXS data for mP13 ZnF5-WWE1-WWE2 is shown in the top plot as log(intensity) versus scattering angle (q). In the lower plots, theoretical scattering for the indicated models is overlaid on the experimental data. Chi^2^ values represent the fit between the experimental and theoretical scattering. Right column. The top plot shows the probability distribution of interatomic distances, P(r), as calculated for the experimental data. The lower plots show the theoretical distribution for the indicated models, overlaid on the experimental data. The DMAX (blue text) and RG (brown text) values obtained from the probability distribution for each plot are denoted. (C) The AlphaFold prediction for mP13 ZnF5-WWE1-WWE2 aligned to the compact monomer model based on the crystal structure. AlphaFold is colored dark gray, except for the ZnF5 region in dark red, and two regions of low confidence modeling that are shown in green (one near the WWE2 binding site, and one in the linker connecting WWE1 and WWE2). The compact monomer is colored light gray, except for the partially ordered ZnF5 in red, and the regions corresponding to the low confidence AlphaFold predictions in blue. The inset focuses on the ZnF5 region in greater detail. (D) Electron density distribution resulting from DENSS analysis of mP13 scattering data, with the map colors representing regions of decreasing electron density: red (most dense), orange, yellow, green, and blue (least dense). The compact monomer model based on the crystal structure was aligned to the map by DENSS. (E) Electron density distribution resulting from DENSS analysis of hP13 scattering data, with map colors as in panel D. The AlphaFold model for hP13 ZnF5-WWE1-WWE2 was aligned to the map by DENSS.

We further analyzed the solution properties of mP13 and hP13 using SEC coupled with small-angle X-ray scattering (SEC-SAXS) (Figure 2B and Supplemental Figure 1A,B,C,D), which can provide molecular mass and size information as well as the overall solution conformation. Buffer-subtracted SAXS profiles were processed to obtain structural parameters such as radius of gyration (R_G_), maximum particle dimension (D_MAX_), and molecular weight (Table 2). Consistent with the SEC-MALS results, mP13 and hP13 primarily existed as monomers in solution based on SAXS measurements. The experimental SAXS data for mP13 ZnF5-WWE1-WWE2 was first compared to theoretical scattering information calculated for the following structures: the crystal monomer, the crystal dimer, and the modeled compact monomer (Figure 2B). The theoretical scattering of the crystallographic dimer was an obvious mismatch to the experimental scattering with a poor chi^2^ value of 2.69 and a substantially different R_G_ (Table 2). Moreover, the pair distribution function, P(r), of the crystal dimer was skewed toward larger interatomic distances than observed in the experimental data. The crystal monomer had a chi^2^ value of 2.27, but still exhibited deviations from the experimental scattering data. For example, the crystal monomer exhibits a substantially bi-modal P(r) distribution that reflects the separated WWE domains, but the bi-modal distribution is not evident in the experimental data. The compact monomer had a chi^2^ value of 2.05 and was thus a reasonable match to the experimental scattering, but shorter interatomic distances were overrepresented in the P(r) distribution. We reasoned that the partially disordered ZnF5 region in our compact monomer model led to an underestimation of longer interatomic distances. We extracted the ZnF5-WWE1-WWE2 region of mP13 from the AlphaFold database entry Q3UPF5 in order to obtain a structural prediction for a fully folded ZnF5. There was good overall agreement between the AlphaFold model and the compact monomer (Figure 2C), with the largest difference related to the ZnF5 region, as expected. The AlphaFold model predicted a standard CCCH-type zinc finger fold that extended further away from the WWE1 fold than what we observed with our partially structured ZnF5. The AlphaFold model had a chi^2^ value of 1.21 and was also the best match to the P(r) distribution, with an increased proportion of longer interatomic distances that reflect the fully formed ZnF5 fold (Figure 1B).

**Table 2.** SAXS-based solution structure parameters.

|  | ZnF5-WWE1-WWE2 |  |
| --- | --- | --- |
|  | mouse PARP13 | human PARP13 |
| **Radius of gyration, $R_G$ (Å)** | 20.9 | 20.2 |
| **Maximum dimension, $D_{MAX}$ (Å)** | ~70 | ~65 |
| **Molecular weight (kDa)** | 21.3 | 19.8 |

$R_G$ values were determined using Guinier analysis, $D_{MAX}$ values were estimated using GNOM (Svergun, 1992), and molecular weights were calculated using the $V_C$ method (Rambo and Tainer, 2013), as implemented in RAW (Hopkins, Gillilan and Skou, 2017).

Comparison of the AlphaFold prediction and our compact monomer of mP13 ZnF5-WWE1-WWE2 highlights the relationship between the partial ZnF5 structure and the predicted ZnF5 structure (Figure 2C). Zinc-coordinating residue H505 is in a roughly similar position in both structures, held near one end of WWE1. In the crystal structure, residues C488 and C496 are located on two anti-parallel beta-strands on the side of WWE1 and positioned away from H505; C501 was located in the disordered region that was not modeled. In the AlphaFold model, C488, C496, and C501 are clustered around H505 to form the zinc(II) binding site. A transition between the two models would require a re-structuring of the beta strands bearing C488/C496 to form a helical structure. Given that zinc content analysis indicated approximately 97% saturation with zinc (Figure 1B), we were surprised that there was no bound zinc in our structure and that ZnF5 was only partially structured. The beta-strand bearing C488 interacts with the extended linker region (Figure 1D), suggesting that the formation of the dimer in the crystal lattice might have driven the destabilization of ZnF5.

We also used *ab initio* modeling to further assess the solution conformation of mP13 ZnF5-WWE1-WWE2. Using the mP13 experimental SAXS profile, the electron density distribution was modeled using DENSS, which reported a Fourier coefficient resolution of 25.5 Å. Consistent with the scattering profile and P(r) distribution analysis, the crystal dimer and the crystal monomer were poor matches to the shape of the DENSS map, due to the overall size of the dimer and the separation of the WWE domains in the monomer. The compact monomer of mP13 ZnF5-WWE1-WWE2 was a good match to the shape of the DENSS map, except for a region of additional density that was not occupied by the compact monomer (Figure 1D). The compact monomer of mP13 ZnF5-WWE1-WWE2 was automatically aligned to the DENSS map, which positioned the extra density next to the partially structured ZnF5 in our compact monomer model (Figure 1D). We thus interpreted the additional density to represent the fully folded ZnF5 as predicted in the AlphaFold model. Indeed, we also analyzed hP13 ZnF5-WWE1-WWE2 by SEC-SAXS and observed good correspondence between the experimental scattering curve and the theoretical scattering of the AlphaFold prediction (Q7Z2W4; Supplemental Figure 1C). Furthermore, the human PARP13 AlphaFold prediction matched well to the shape of the DENSS map, including the region corresponding to the additional density (Figure 1E). We obtained similar results using the program DAMMIF to generate bead models representing the scattering data (Supplemental Figure 1E,F).

Thus, the solution-based SAXS analysis is most consistent with a compact monomer conformation for ZnF5-WWE1-WWE2, as seen with the good agreement between the AlphaFold prediction and the experimental SAXS data. Our crystal structure indeed captured the essential attributes of the compact conformation of ZnF5-WWE1-WWE2, albeit arising from two different protein chains. We thus view the crystal structure as representative of the monomeric conformation of ZnF5-WWE1-WWE2 and appropriate for analyzing ADPr/ATP ligand interactions, which are not predicted by AlphaFold. Indeed, one of the low-confidence regions of the AlphaFold prediction abuts the binding cavity of WWE2 (Figure 2C).

### Variation in the WWE domain structure and binding properties

Although mP13 ZnF5-WWE1-WWE2 was crystallized in the presence of 1 mM ADPr or 1 mM ATP, ligand was only observed bound to WWE2 (Figure 3A,B). There was no evidence for ligand interaction with WWE1, despite the domain being accessible within the crystal lattice. The binding pocket of WWE2 is formed at one end of a barrel-shaped fold composed of six β-strands and one a-helix (Figure 3A,B). The loops connecting the secondary structure elements and the central cavity at one end of the barrel contribute the major interacting residues (Figure 3C), centered around W585 that engages one face of the adenine base (W611 in hP13). Specific contacts are also made with the adenosine ribose and two phosphate groups of ADPr/ATP; however, there are fewer contacts made with the terminal ribose of ADPr and the terminal *γ*-phosphate of ATP.

**Figure 3:**
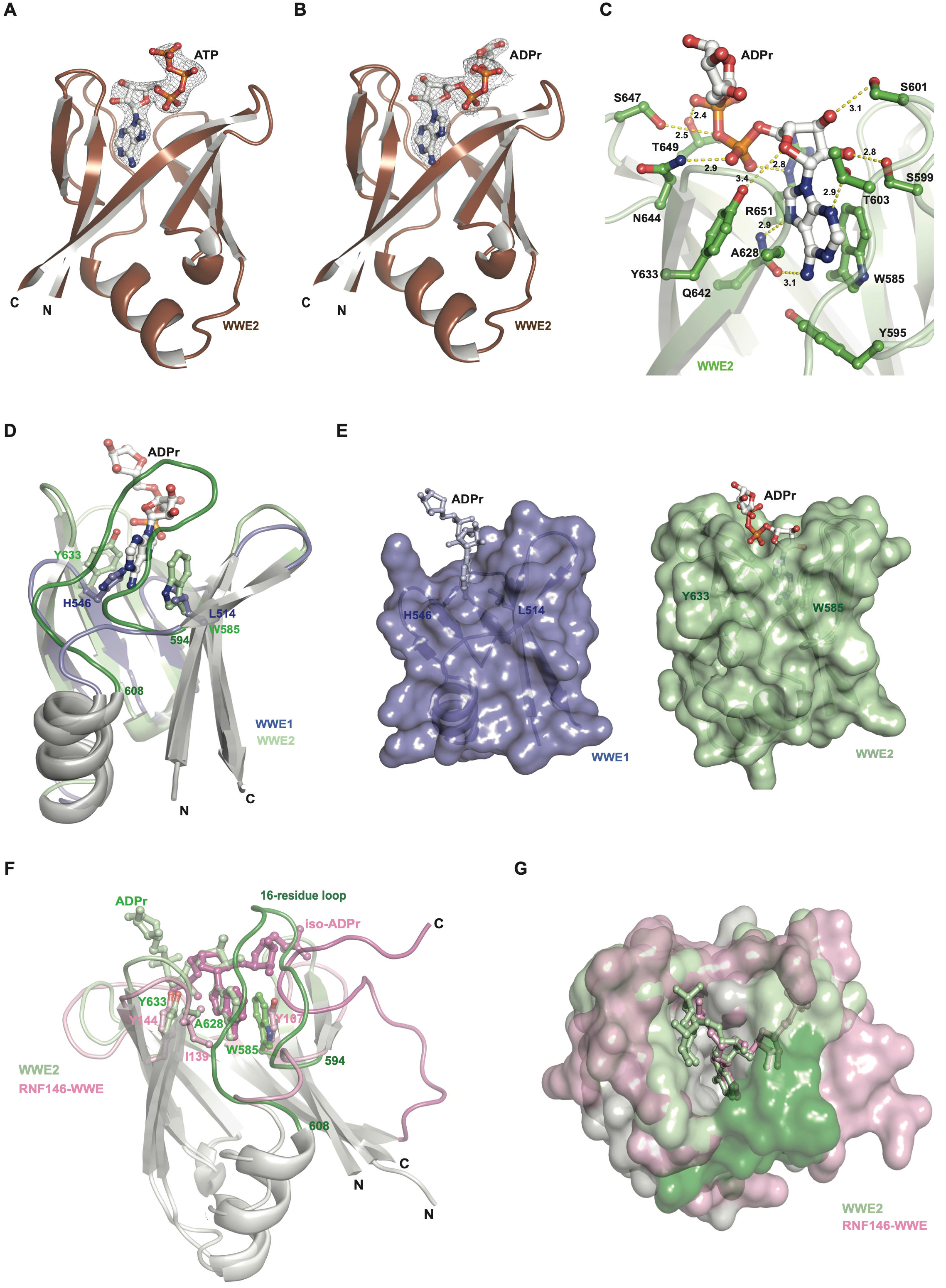
Ligand-binding properties of the WWE2 domain. (A) ATP bound within the cavity of WWE2. A weighted 2F_O_-F_C_ electron density map is shown around the ATP molecule and is contoured to the 1.5σ level. (B) ADPr bound within the cavity of WWE2. A weighted 2F_O_-F_C_ electron density map is shown around the ADPr molecule and is contoured to the 1σ level. (C) A view of key WWE2 residues interacting with ADPr. Interatomic distances (Å) are indicated next to dashed lines connecting certain atoms. (D) Structure of WWE1 (blue) superimposed on WWE2/ADPr of mP13 (green). (E) Surface representation of WWE1 (on left in blue) illustrating the lack of a binding cavity compared to WWE2 (on right in green) with a deep cleft forming the ADPr binding pocket. (F) Structure of the WWE domain of RNF146 bound to iso-ADPr (PDB code 3V3L) in light pink superimposed on WWE2 of mP13 bound to ADPr (green). (G) Surface representation of WWE from RNF146 bound to iso-ADPr (light pink) superimposed on WWE2 of mP13 bound to ADPr (green).

A comparison of the WWE1 and WWE2 domains provides a clear structural basis for the observed differences in binding to ligand (Figure 3D,E). WWE1 and WWE2 share the same barrel-shaped fold, but the connecting loops are much shorter in WWE1. Most notably, one loop present in WWE2 (residues 594 to 608) is over 10 residues shorter in WWE1, entirely removing a major portion of the binding surface. The cumulative effect of the smaller WWE1 is that the binding pocket is much shallower. Indeed, the shape of WWE1 does not resemble a binding pocket (Figure 3E). There is also an offset in the β-strands of WWE1 relative to WWE2 that displaces key ligand-binding residues (Figure 3D). Thus, the WWE1 domain lacks essentially all the WWE2 properties responsible for interaction with ADPr/ATP.

Our initial work indicated that hP13 and mP13 bind to ATP and ADPr, and we thus used these ligands to aid in crystallization efforts and to provide insights into ZnF5-WWE1-WWE2 binding properties. However, we noted that other WWE domains, such as the well-studied WWE domain from RNF-146, do not bind ADPr (Wang *et al*., 2012). Rather, RNF-146 is capable of binding iso-ADPr, a molecule in which the adenosine monophosphate ribose is connected to a ribose phosphate (Figure 3F), thus modeling the primary ribose-ribose (2′-1″) linkage of poly(ADP-ribose) (18). A comparison of the RNF-146 WWE/iso-ADPr structure (PDB code 3V3L) (18) and the mP13 WWE2/ADPr structure revealed important differences that are likely to explain the distinct binding properties (Figure 3F,G).

Adenosine monophosphate occupies similar positions in both structures, with the adenine base sandwiched between hydrophobic residues W585, Y633 and A628 in WWE2, and Y107, Y144 and I139 in RNF146 (Figure 3F). A major difference is a 16-residue loop in WWE2, comprised of residues Q594 to N608, that is entirely missing in RNF-146. Instead, RNF-146 has a C-terminal loop that forms part of the binding site for the ribose phosphate group. The16-residue loop of WWE2 extends further into the binding cavity to contribute residues that directly engage the adenosine ribose, and in this position the loop sterically prohibits the possibility of a second ribose group (Figure 3F,G). The 16-residue loop therefore changes the profile of the WWE2 cavity for ligand binding, forming a narrower opening than RNF-146, which is widened to accept the second ribose group (Figure 3G). Hence, RNF-146 appears to be dependent on the second ribose phosphate to establish efficient binding. In contrast, WWE2 appears tailored to recognize a terminal adenosine ribose without a ribose extension, and in fact would appear to disfavor the ribose-ribose linkage, without some form of adaptation in the positioning of the 16-residue loop.

We directly measured the binding capacity of hP13 ZnF5-WWE1-WWE2 with a fluorescence polarization (FP) assay using ATP labeled with fluorescein on the g-phosphate (ATP-FAM), as the *γ*-phosphate was largely free of contacts in the ZnF5-WWE1-WWE2/ATP structure. hP13 ZnF5-WWE1-WWE2 interacted with the ATP-FAM with an apparent binding affinity of 218 ±19 nM (Figure 4A). In contrast, RNF-146 showed no real evidence of binding to ATP-FAM in our fluorescence polarization assay (Figure 4A), consistent with a published report (Wang *et al*., 2012). We also tested whether we could abolish the binding activity of WWE2 through mutagenesis, in which we targeted the central W611 in the binding cavity. The mutant W611A of hP13 ZnF5-WWE1-WWE2 showed no evidence of binding to ATP-FAM, consistent with WWE2 representing the primary binding site. We also assessed binding using a DSF assay to monitor relative thermal stability in the absence/presence of ligand, with the expectation that interaction with ligand will increase thermal stability (Figure 4B). We indeed observed an increase in the thermal stability of hP13 ZnF5-WWE1-WWE2 in the presence of ATP and ADPr. In contrast, the W611A mutant did not show an increase in thermal stability in the presence of ATP/ADPr. The W611A mutant exhibited a decrease in thermal stability relative to wild-type, suggesting that the mutation in the central cavity has somewhat altered protein stability. However, we observed no problems in overexpressing and purifying this mutant, suggesting that the overall fold is intact.

**Figure 4:**
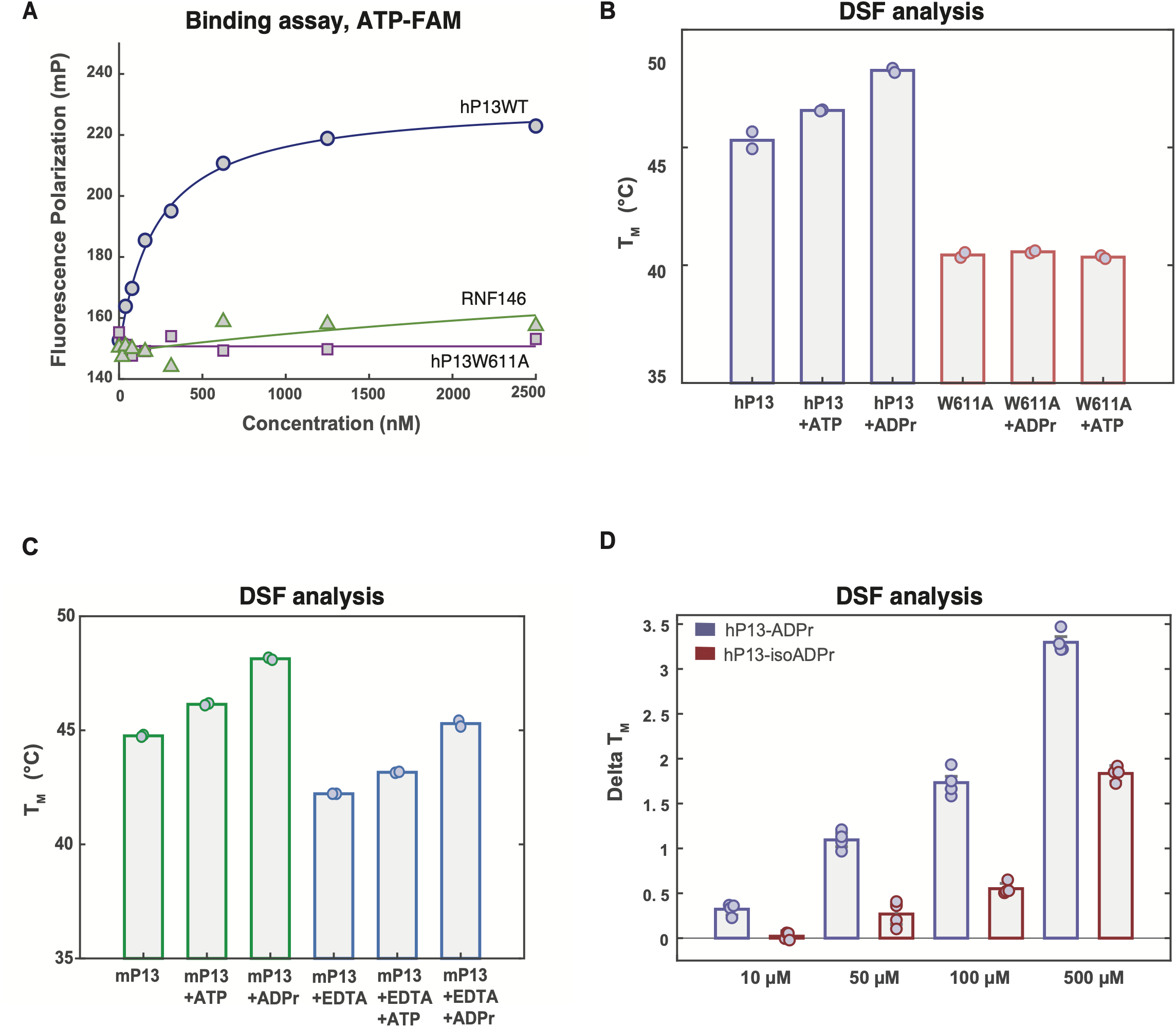
Biophysical and biochemical analysis of mP13 and hP13 ZnF5-WWE1-WWE2. (A) Representative curves from the FP binding assay using ATP-FAM and the indicated proteins. A 1:1 binding model was fit to the data (solid lines). These experiments were repeated at least three times to provide average K_D_ values and associated standard deviations. (B) DSF analysis of relative thermal stability for the indicated proteins in the absence of ligand or in the presence of ATP or the presence of ADPr. (C) DSF analysis comparing untreated (green bars) and EDTA-treated mP13 ZnF5-WWE1-WWE2 (blue bars) in the absence of ligand or with ATP or with ADPr. (D) ΔTm comparison from DSF analysis performed with hP13 ZnF5-WWE1-WWE2 and four concentrations of ADPr or iso-ADPr.

mP13 ZnF5-WWE1-WWE2 also showed evidence of interaction with ATP and ADPr using DSF (Figure 4C). Our structure of mP13 ZnF5-WWE1-WWE2 with a partially structured ZnF5 was still capable of interaction with ADPr, suggesting that ZnF5 does not directly contribute to ADPr/ATP binding. We tested biochemically whether loss of bound zinc, and thus ZnF5 structure, influenced ligand binding by treating ZnF5-WWE1-WWE2 with 10 mM EDTA to leech away zinc. The treated sample exhibited a decrease in thermal stability relative to a mock-treated sample (Figure 4C), and we interpret the decrease in thermal stability as reflecting a loss of zinc from ZnF5. The treated sample still showed evidence of interaction with ATP and ADPr (Figure 4C). Collectively, the data support the conclusion that the WWE2 domain serves as the primary interaction site with ADPr and ATP, and that ZnF5 and WWE1 do not contribute to this binding activity.

We also used DSF to evaluate hP13 ZnF5-WWE1-WWE2 interaction with ADPr compared to iso-ADPr. Based on structural comparison (Figure 3F,G), iso-ADPr was predicted to be a poor fit to the binding pocket of WWE2, due to a 16-residue loop blocking the extension of the second ribose of iso-ADPr. Over a series of concentrations, we observed again that ADPr increased hP13 ZnF5-WWE1-WWE2 thermal stability (Figure 4D). We also observed an increase in thermal stability in the presence of iso-ADPr, but to a lesser extent than that seen with ADPr, and the change in thermal stability required higher concentrations of iso-ADPr (Figure 4D). The DSF results suggest a more robust interaction with ADPr, but that iso-ADPr is nonetheless capable of forming an interaction. We have not been able to co-crystallize ZnF5-WWE1-WWE2 with iso-ADPr, which could reflect that the interaction is unfavorable. We infer that the 16-residue loop is capable to some extent of adapting to the structure of iso-ADPr, but that the WWE2 binding cavity favors ADPr.

### PAR binding properties of ZnF5-WWE1-WWE2

We first analyzed PAR binding through an FP-based competition assay. The relative affinity of different ligands was assessed by their capacity to outcompete ATP-FAM binding to hP13 ZnF5-WWE1-WWE2, with the expectation that an efficient competitor would lower the FP signal (Figure 5A). The ATP-FAM concentration was 5 nM, and hP13 ZnF5-WWE1-WWE2 was used at 100 nM to provide a substantial population of bound ATP-FAM (roughly 50%) and thus a stable FP measurement. PAR chains of 3 units and 6 units (3-mer and 6-mer) were compared to ADPr, representing a single PAR unit. ADPr behaved in large part like a direct competitor of the ATP binding site, as expected, given that the interactions with ATP and ADPr were virtually identical in our structures. 3-mer PAR behaved similarly to ADPr, in that the FP signal was lowered to a comparable extent as the concentration increased. Of note, difficulties in producing PAR of defined lengths in great amounts limited the concentrations that could be tested in this assay. The similarity in the results using ADPr and 3-mer PAR suggested that the terminal end of the PAR chain was competing for the WWE2 binding cavity, and that the presence of 3 PAR units did not increase the capacity to compete for binding. In contrast, 6-mer PAR was observed to be a much better competitor ligand, with a substantial reduction in FP signal evident at the first concentration tested. Given the different capacities of 3-mer and 6-mer to serve as competitors, we inferred that PAR chains longer than a certain minimal length have the capacity to engage other parts of the hP13 ZnF5-WWE1-WWE2 structure and thereby gain a competitive advantage. As presented later, structural analysis indicates a prominent groove that could engage the PAR chain.

**Figure 5:**
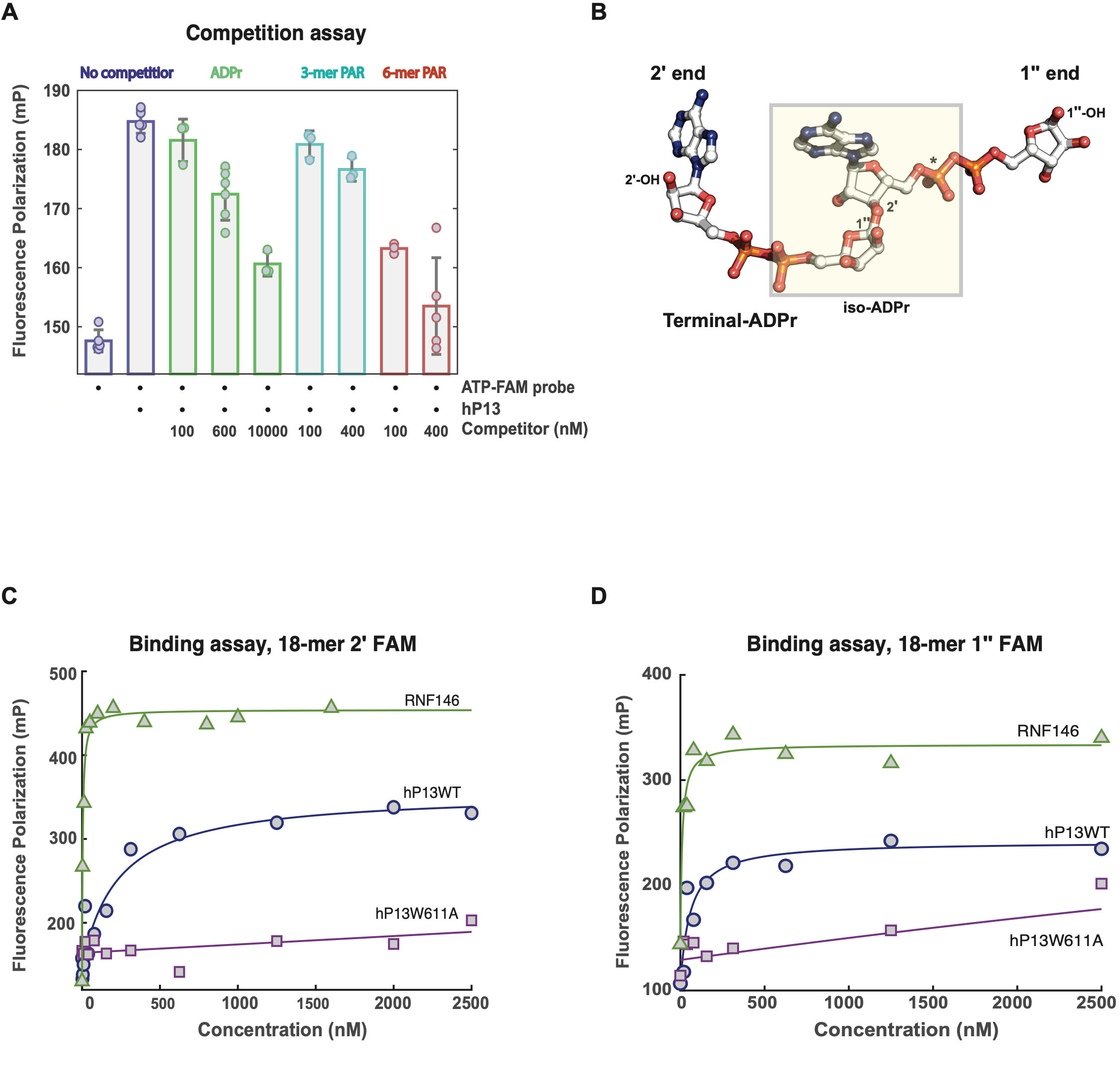
PAR binding properties of ZnF5-WWE1-WWE2. (A) FP competition assay comparing ADPr, 3-mer PAR, and 6-mer PAR capacity to outcompete ATP-FAM probe binding to hP13 ZnF5-WWE1-WWE2. (B) A short PAR structure of two units illustrates the two termini of the PAR chain. The iso-ADPr unit is highlighted in pale yellow. (C and D) Representative curves from PAR binding analysis using 18-mer PAR labeled on the 1″ end (panel C) or using 18-mer PAR labeled on the 2′ end (panel D). A 1:1 binding model was fit to the data (solid lines). The experiments were repeated at least three times to produce average K_D_ values and the associated standard deviations.

We directly measured the PAR-binding capacity of hP13 ZnF5-WWE1-WWE2 using fluorescently labeled versions of an 18-mer PAR chain in FP binding assays. The ELTA method provides a convenient and robust strategy to label the terminal adenosine group of a PAR chain with dATP (Ando *et al*., 2019), thus we added a dAMP-fluorescein group to the 2′ terminus (Figure 5B) of a purified 18-mer PAR molecule (18-mer 2′ FAM). However, given our model of WWE2 making direct contacts with the terminal adenosine group of a PAR chain, we also created 18-mer PAR labeled on the opposite 1″ end of the chain (Figure 5B) with fluorescein (18-mer 1″ FAM). Labeling of the 1″ end takes advantage of a terminal adenosine phosphate that remains after PAR is released from proteins using base treatment. hP13 ZnF5-WWE1-WWE2 was estimated to bind 18-mer 1″ FAM with a K_D_ of 68 ± 34 nM. This K_D_ value is roughly 3-fold lower than the K_D_ obtained for hP13 ZnF5-WWE1-WWE2 binding to ATP-FAM, consistent with the hypothesis that PAR chains provide an additional mode of interaction. hP13 ZnF5-WWE1-WWE2 was estimated to bind 18-mer 2′ FAM with a K_D_ of 278 ± 64 nM (Figure 5D), thus with 4-fold lower affinity than observed with 18-mer 1″ FAM. We interpret this difference in affinity to reflect that the dAMP-fluorescein from ELTA labeling on the 2′ end adenosine group has created conflict with the 16-residue loop of WWE2 that forms direct contacts with the adenosine ribose (Figure 3C,D). Notably, RNF-146 bound to both 18-mer PAR chains with similar K_D_ values of 3.81 ± 0.72 nM and 4.78 ± 0.97 nM (Figure 5C,D), consistent with the observation that RNF-146 does not specifically engage the terminal adenosine ribose. The W611A mutant of hP13 ZnF5-WWE1-WWE2 showed no evidence of stable interaction with either of the 18-mer PAR molecules (Figure 5C,D), suggesting that WWE2 represents the dominant site of interaction with PAR.

### Structural evidence for an extended PAR binding site

Inspection of the shape and electrostatic properties of mP13 ZnF5-WWE1-WWE2 suggested a surface groove that could function as an extended PAR binding site (Figure 6A), as suggested by the binding studies. Since the mP13 ZnF5-WWE1-WWE2 structure exhibited a partially folded ZnF5, we analyzed our structure with and without this region included in the surface representation and electrostatic surface potential (Figure 6A,B), and we also performed the analysis with the AlphaFold model (Figure 6C). Extending from the ADPr/ATP binding cavity of WWE2 in each of the models, a positively charged groove is traced along the surface of WWE2 and continues to a deepened region of the groove at the junction of WWE2 and WWE1. The deepened region of the surface groove exhibits a high density of positive surface charge, and we anticipate that this site represents a major point of contact with PAR chains that extend from an anchor point in WWE2 (Figure 6A). It is compelling that the distance between the WWE2 binding site and the deep region of the surface groove appears to set a minimal length requirement for PAR chains to engage both sites, consistent with our evaluation of 3-mer and 6-mer PAR as competitor ligands. The size of the main pocket within the groove varies based on the modeled position of ZnF5, so it is possible that ZnF5 will be situated to have an impact on this putative binding region. Sequence analysis of the ZnF5-WWE1-WWE2 region from multiple species indicates strong conservation within the groove along the outside of the WWE1 fold, whereas in contrast the non-functional binding cavity of WWE1 exhibited low sequence conservation (Supplementary Figure 2). We can thus speculate that the function of the WWE1 fold is to contribute surface residues to form the extended groove, rather than contributing a functional binding cavity as seen in WWE2. We also note that in each of the mP13 crystal structures determined, a phosphate ion was modeled at the edge of the deep groove, suggesting a possible binding site for additional ADPr units (Figure 6D). In particular, conserved residue R656 engages the phosphate ion. Taken together, we propose that ZnF5-WWE1-WWE2 anchors preferentially on the terminal ends of PAR chains using the WWE2 binding cavity, and PAR chains of at least 5 or 6 units will have the capacity to form a more stable interaction by engaging the surface groove (Figure 6A).

**Figure 6:**
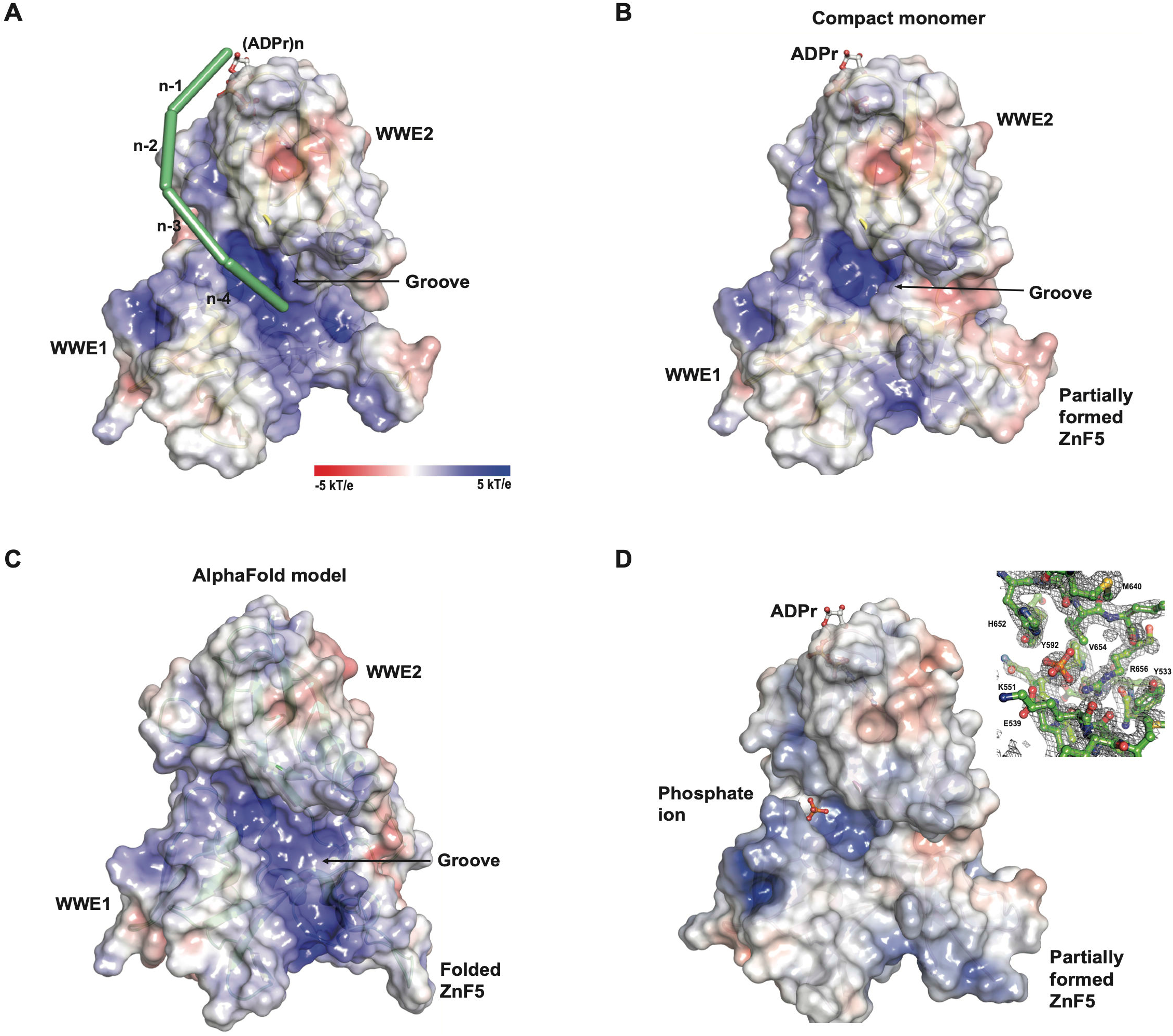
A putative PAR binding groove. (A) Electrostatic surface potential of the mP13 compact monomer with the ZnF5 region removed, revealing a positively charged groove extending from the ADPr binding cavity in WWE2. The putative binding groove is highlighted with units of PAR, represented as green bars, tracing along the groove on the surface of WWE1 and WWE2. The terminal ADPr in the binding pocket is designated n, and the preceding ADPr units are designated n-1, n-2, and so forth, providing a rough estimate of the distance between the WWE2 binding pocket and the deepened region of the groove. (B) Electrostatic surface potential of the mP13 compact monomer with the partially formed ZnF5 included. (C) Electrostatic surface potential of the AlphaFold model. (D) Electrostatic surface potential of the compact monomer, highlighting the position of a bound phosphate ion within the groove. The inset panel shows the atomic model surrounding the phosphate ion, with the electron density map for the region overlaid (weighted 2F_O_-F_C_ contoured at 1.5σ).

## DISCUSSION

Our crystal structure of mP13 ZnF5-WWE1-WWE2 indicated different possibilities for the relative arrangement of these domains. Solution-based biophysical analysis indicated that the compact, monomeric arrangement of ZnF5-WWE1-WWE2 is most representative of the structure of this fragment. We note that an AlphaFold model predicts a similar compact arrangement of domains, and a study in BioRxiv reports a structure of hP13 ZnF5-WWE1-WWE2 that also appears to adopt the compact, monomeric conformation (structure coordinates not available; (Xue *et al*., 2020)). However, the SEC-MALS analysis did suggest that a small population of ZnF5-WWE1-WWE2 dimer exists in solution. The abundance of the dimer in solution appeared to correlate with the abundance of ZnF5-WWE1-WWE2 that was not bound to zinc, suggesting that the conformation of ZnF5-WWE1-WWE2 could be regulated by zinc availability and that ZnF5 could be a dynamic regulator rather than a static structural element.

The dimer of ZnF5-WWE1-WWE2 observed in the crystal structure could represent a conformation adopted under certain conditions, for example when P13 acts to signal the detection of viral RNA. In the dimer conformation, the WWE domains form the same interfaces seen in the compact monomer conformation, but they originate from different polypeptides. The major new contacts in the dimer conformation involve the linker region of both molecules. The linker regions form an extended interface in the dimer, and in the compact monomer the linker folds back onto itself. Based on sequence analysis, both ends of the linker region are well conserved, but the center of the linker is only moderately conserved across species (Supplementary Figure 2). It could be that the inherent flexibility of the linker allows the WWE domains to exist in alternate conformations to perform diverse functions. Further studies are required to establish the relevance of the dimer conformation and the potential connection to PARP13 cellular function.

Our co-crystal structures with ATP and ADPr have provided the first views of ligands bound to ZnF5-WWE1-WWE2. The central cavity of WWE2 and surrounding loops form the binding site that primarily engages the adenosine group of ATP and ADPr. The WWE2 binding site is distinct from the WWE domain of RNF-146, which is tailored to recognize the ribose-ribose linkage of poly(ADP-ribose) (Wang *et al*., 2012). Rather, WWE2 engages the ribose group of adenosine in a manner that should disfavor a ribose extension. WWE1 lacks entirely the central binding cavity, and the structure appears to contribute structurally to the extended PAR binding site that we have inferred through binding analysis and that is supported by structural analysis (Figure 6).

Collectively, our analysis supports a PAR binding site that extends beyond the WWE2 cavity observed in the crystal structure. The extended PAR binding site was evident in the competition assay in which the 6-mer PAR molecule was a much better competitor against ATP-FAM, compared to 3-mer PAR and ADPr provided at the same concentrations. Labeling of 18-mer PAR on the terminal ADPr group using the ELTA method introduces a fluorescein-dAMP group that would interfere with the narrow pocket of the WWE2 cavity. Indeed, 18-mer PAR labeled in this manner exhibited 4-fold weaker binding affinity than 18-mer PAR labeled on the other end (70 nM versus 280 nM). For the 18-mer PAR bearing the obstruction on the terminal ADPr, we attribute the 280 nM binding affinity to at least partially arise from the extended binding groove on ZnF5-WWE1-WWE2.

RNF-146 recognition of the ribose-ribose linkage was not influenced by differences in the label location on 18-mer PAR, since the ribose-ribose linkage is the PAR feature recognized with its WWE domain, rather than the terminal adenosine group. ZnF5-WWE1-WWE2 thus represents a new mode of engaging poly(ADP-ribose) that is sensitive to the structure of the terminal end of the chain, and to the length of the chain. These features could tune PARP13 cellular function to be responsive to modifications of poly(ADP-ribose) structure, including modification of the terminal end by enzymes such as OAS1, or changes in chain length through the action of poly(ADP-ribose) degrading enzymes. The dependency on the length of the PAR chain as a means to both enhance binding affinity and extend the binding surface further than the ADPr binding pocket has been noted in studies performed on the macrodomain of ALC1 (Singh *et al*., 2017).

In summary, our study provides new structural and functional insights into the PAR binding properties of PARP13/ZAP. The structure and binding studies provide new avenues for investigating the cellular functions of PARP13/ZAP, including the contribution of PAR binding and potentially aspects regulating conformational changes in response to the detection of viral RNA.

## MATERIALS AND METHODS

### Gene cloning and mutagenesis

hP13 ZnF5-WWE1-WWE2 (residues 507 to 699) was cloned as a His-tagged, GB1 (B1 domain of Streptococcal protein G) fusion protein with a TEV protease cleavage site in a modified pET28 vector. A codon-optimized gene for mP13 ZnF5-WWE1-WWE2 (residues 476 to 673) was cloned as a His-tagged, SMT (SUMO-like tag) fusion protein in a pET28a vector (Genscript). Quick Change mutagenesis was performed to generate the W611A mutant of hP13 ZnF5-WWE1-WWE2, which was confirmed by automated Sanger DNA sequencing. The WWE domain of RNF-146 was produced as fusion with GST as described (DaRosa *et al*., 2015).

### Protein expression and purification

mP13 ZnF5-WWE1-WWE2 and hP13 ZnF5-WWE1-WWE2 (wild-type and mutant W611A) were expressed in Rosetta2 *E. coli* cells using LB medium and IPTG induction (0.2 mM) at 16°C for 18 hours. Seleno-methionine mP13 ZnF5-WWE1-WWE2 was expressed in defined media as described. Pelleted cells were resuspended in the following buffer: 25 mM HEPES pH 8.0, 500 mM NaCl, 0.1 mM TCEP. mP13 ZnF5-WWE1-WWE2 (wild-type and seleno-methionine) and hP13 ZnF5-WWE1-WWE2 (wild-type and mutant W611A) were purified using Ni^2+^ affinity chromatography on a 5 mL His-trap column, gel filtration chromatography on either a Sephacryl 75 or Sepharose 200 column, and ion exchange chromatography using a 5 mL Q-Sepharose column (Cytiva). After the Ni^2+^ column, proteins were dialysed overnight at 4°C to remove 400 mM imidazole that was used for elution (25 mM HEPES pH 8.0, 300 mM NaCl, 0.1 mM TCEP). The SMT or GB1 tags were cleaved by addition of ULP1 or TEV protease, respectively, during dialysis. The cleaved and dialysed proteins were passed over a 5 mL His-trap column to remove the his-tagged SMT, GB1, and proteases. The flow-through was concentrated by centrifugation and loaded for gel filtration chromatography (Sephacryl 75 for hP13 ZnF5-WWE1-WWE2, and Sepharose 200 for mP13 ZnF5-WWE1-WWE2). Gel filtration fractions containing proteins of interest were passed over a Q-Sepharose column, in which the protein of interest flowed through, and remaining contaminants bound to the column. The flow-through of the Q-Sepharose column was concentrated by centrifugation and flash-frozen in small aliquots in liquid nitrogen and stored at −80°C.

### Crystallization and X-ray structure determination

mP13 ZnF5-WWE1-WWE2 (0.4 mM) was incubated with 1 mM ATP or with 1 mM ADP-ribose (ADPr) prior to setting up crystallization plates at room temperature. Crystals appeared in 1.4 M Na/K phosphate pH 5.5 within two to three days with either ATP or ADPr. Crystals were cryo-protected by supplementing growth conditions with 30% glycerol or 30% ethylene glycol prior to flash-cooling in liquid nitrogen for diffraction experiments. X-ray diffraction data for mP13 ZnF5-WWE1-WWE2 crystals with ADPr and with ATP were collected at the CMCF-BM (08B1-1) beamline at the Canadian Light Source and the data were processed using XDS (Kabsch, 2010). Seleno-methionine containing crystals of mP13 ZnF5-WWE1-WWE2 were obtained in the same conditions as above. Single-wavelength anomalous dispersion (SAD) X-ray diffraction data from a single crystal grown with ATP was collected at the Advance Light Source beamline 8.3.1 and processed using XDS (Kabsch, 2010). PHENIX AUTOSOL and AUTOBUILD were used to determine experimental phases and to partially build the structure (Adams *et al*., 2010). The seleno-methionine structure was then used as the starting model for the wild-type ZnF5-WWE1-WWE2 complexes with ATP and with ADPr. Manual model building was performed using COOT (Emsley *et al*., 2010) and model refinement was carried out using PHENIX.

### SEC-MALS

Size-exclusion chromatography (SEC) was performed on an ÄKTAmicro liquid chromatography system (Cytiva). Samples were injected onto a pre-equilibrated Superdex 200 Increase 10/300 GL column operated at a flow rate of 0.35 ml/min with the following buffer: 25 mM HEPES pH 8.0, 150 mM NaCl, 0.1 mM DTT, and 1 mM EDTA. Samples then flowed in-line through a multi-angle light scattering (MALS) detector (DAWN HELEOS II, Wyatt Technology) followed by a refractive index detector (OptiLab T-rEX, Wyatt Technology). Data were analyzed with ASTRA 6.1.6.5 software (Wyatt Technology) to determine the molecular weight and the percent mass fractions of the eluting peaks. BSA was used as a standard for data collection and analysis prior to injecting the samples of interest.

### SEC-SAXS

For SEC-SAXS, an ÄKTAmicro FPLC (Cytiva) was connected to a BioXolver SAXS system (Xenocs) equipped with a MetalJet D2+ 70 kV X-ray source (Excillum) and a PILATUS3 R 300K detector (Dectris). The samples were injected at 100 mg/mL for mP13 ZnF5-WWE1-WWE2 and at 17 mg/mL for hP13 ZnF5-WWE1-WWE2 onto a pre-equilibrated Superdex 200 Increase 10/300 GL column (25 mM HEPES pH 8.0, 150 mM NaCl, 0.1 mM DTT, and 1 mM EDTA) operated at 0.1 mL/min. X-ray scattering data was collected in 30-second exposures (~400 total exposures) over the course of the elution profile. Average scattering of a buffer only region was subtracted from the average scattering over a protein peak to yield the buffer-subtracted scattering profile. The radius of gyration (R_G_), forward scattering intensity (I_0_), and maximum dimension of (D_MAX_) were derived using RAW software version 2.1.0 (Hopkins, Gillilan and Skou, 2017). FoxS and CRYSOL (ATSAS software version 2.8.4) was used to perform comparisons of experimental SAXS profiles to various models (Svergun, Barberato and Koch, 1995; Schneidman-Duhovny *et al*., 2013). RAW was also used to perform DAMMIF/N and DENSS reconstructions for the SAXS profile of both mouse and human samples (Franke and Svergun, 2009; Grant, 2018).

### Zinc content analysis

Protein samples at 10 μM were dialysed overnight at 4°C against 20 mM Hepes pH 8.0, 150 mM NaCl, 0.1mM TCEP, and 1 mM EDTA. A Perkin Elmer NexION 300x was used for the quantification of zinc. ICP-MS standards certified traceable to NIST were used. All standards and blanks were prepared using ultrapure water (Milli-Q, 18.2 MΩ cm; total organic carbon <1 μg L^-1^) and ultra-trace nitric acid (Plasma Pure Plus, SCP Science). EPA 200.7 standard 6 purchased from High-Purity Standards was used for calibration. ICP-MS standards QCP-QCS-3 (Inorganic Ventures) and QCS-27 (High Purity Standards) were used for quality control. Yttrium (Inorganic Ventures) was used as internal standard. The isotope ^66^Zn was monitored. ICP-MS analyses were performed in triplicate measurements with 20 readings each and an integration time of 1 second.

### Differential scanning fluorimetry (DSF)

DSF experiments were performed by adding 8 μM protein to a solution containing 25 mM HEPES pH 8.0, 150 mM NaCl, 1 mM EDTA, 0.1 mM TCEP, and 1x SYPRO-orange dye (Sigma-Alrich), plus or minus the specified ligand concentrations. Iso-ADPr was a kind gift from Dr. Wenqing Xu (University of Washington). Fluorescence emission of the samples was measured on a Roche LightCycler 480 RT-PCR, as the temperature was increased from 20 to 85°C. The Tm values were determined by a Boltzmann distribution fit to the data. The reported ΔT_M_ values represent the T_M_ value with ligand minus the T_M_ value without ligand.

### PAR synthesis and labeling

3-mer, 6-mer, and 18-mer PAR were produced enzymatically and purified using boronate-affinity and ion exchange chromatography (Edwin S. Tan, Kristin A. Krukenberga, 2012; Ando *et al*., 2019). 18-mer PAR was labeled on the 2′ end with fluorescein-dAMP using OAS1 and the ELTA method (Ando *et al*., 2019). The 1″ end was FAM-labeled on the adenosine phosphate that remains after PAR is released from proteins using base treatment (Abraham *et al*., 2020). First, an alkyne-PEG1-amine linker was added to the adenosine phosphate using EDC coupling. The alkyne-PAR was purified to remove excess alkyne-PEG1-amine using the Monarch Nucleic Acid Cleanup Kit (NEB) following the recommended protocol. Second, FAM-PAR was produced through Cu(I)-catalyzed click chemistry between FAM-azide and the alkyne-PAR. Finally, the FAM-PAR was purified via ion-pair reversed-phase high performance liquid chromatography on an Agilent 1260 Infinity II system using an InertSustain C18 HP 3um column (4.6 x 250 mm, GL Sciences) with two mobile phases. Mobile phase A consisted of 100 mM triethylammonium acetate pH 7.5 and mobile phase B consisted of 100% acetonitrile. Fractions containing FAM-PAR were dried down and then resuspended in Milli-Q water to a concentration of 10 μM.

### Fluorescence polarization (FP) binding and competition assays

FP binding affinity measurements were performed using proteins at the designated concentrations in a buffer containing 12 mM HEPES pH 8.0, 25 mM KCl, 50 μg/mL BSA, 4% glycerol, and either 5.5 mM 2-mercaptoethanol or 0.1 mM TCEP. Fluorescent probes (18-mer 1″ fluorescein, 18-mer 2’fluorescein, and ATP-FAM) were included at 5 nM. ATP-FAM [g-(6-Aminohexyl)-ATP-6-FAM] was purchased from Jena Bioscience. Reactions were incubated for 30 minutes at room temperature before taking fluorescence polarization measurements on a Victor3Vplate reader (PerkinElmer). Measurements were buffer subtracted, and the polarization value was calculated for each measurement. A one-to-one binding model was fit to the data with a nonlinear regression equation using Microsoft excel solver to obtain binding constant (K_D_) values. The experiments were performed at least three times, and the averages and standard deviations are reported. FP competition assays were performed in the same buffer as above, using ATP-FAM at 5 nM. ADPr, 3-mer PAR, and 6-mer PAR were included at the designated concentrations to compete with ATP-FAM binding to hP13 ZnF5-WWE1-WWE2. FP readings were taken on a Victor3Vplate reader (PerkinElmer) after an incubation period of 30 minutes. Measurements were buffer subtracted, and the polarization value was calculated for each measurement. The experiments were performed at least three times, and the averages and standard deviations are reported.

## ACKNOWLEDGEMENTS

We thank Dr. Wenqing Xu and Dr. Zhizhi Wang for providing the RNF-146 WWE expression vector and purified iso-ADPr, and Dr. Normand Cyr for assistance in operating the SEC-SAXS instrument. This work was supported by grants from the Canadian Institute of Health Research (PJT153295 to J.M.P) and the National Institutes of Health (R01GM104135 to A.K.L.L). A portion of the work was conducted at the Advanced Light Source (ALS), a national user facility operated by Lawrence Berkeley National Laboratory on behalf of the Department of Energy, Office of Basic Energy Sciences, through the Integrated Diffraction Analysis Technologies (IDAT) program, supported by DOE Office of Biological and Environmental Research. Part of the research described in this paper was performed using beamline CMCF-BM at the Canadian Light Source, a national research facility of the University of Saskatchewan, which is supported by the Canada Foundation for Innovation (CFI), the Natural Sciences and Engineering Research Council (NSERC), the National Research Council (NRC), the Canadian Institutes of Health Research (CIHR), the Government of Saskatchewan, and the University of Saskatchewan. SEC-MALS and SEC-SAXS data were acquired at the Structural Biology Platform of the Université de Montréal, which operates with support from the Canadian Foundation for Innovation (30574).

## AUTHOR CONTRIBUTIONS

J.R.A.K., H.S., and J.M.P conceived the study; J.R.A.K. and H.S. performed protein purification and gene cloning; J.R.A.K. performed crystallography, biophysical analysis, biochemical analysis, and data analysis; M.D., I.U., S-J.C., and A.K.L.L. performed PAR synthesis, labeling, and purification; J.R.A.K. and J.M.P wrote the manuscript with input from all authors; J.M.P. and A.K.L.L directed the research.

## ACCESSION CODES

Coordinates and structure factor amplitudes have been deposited in the Protein Data Bank with the following accession codes: mP13 ZnF5-WWE1-WWE2 with ATP (7SZ2) and mP13 ZnF5-WWE1-WWE2 with ADPr (7SZ3).

---

**Supplementary Figure 1: SEC-SAXS data analysis of PARP13 ZnF5-WWE1-WWE2.** (A) Size exclusion chromatogram of ZnF5-WWE1-WWE2 from mP13 plotted as integrated intensity (left y axis) versus exposure frame number. The RG estimation (right y axis) is shown across the elution peaks (orange boxes). The frames chosen for buffer subtraction are highlighted in pale green. (B) Size exclusion chromatogram of ZnF5-WWE1-WWE2 from mP13 plotted as integrated intensity (left y axis) versus exposure frame number. The molecular mass estimation (right y axis) is shown across the elution peaks (red boxes). The frames chosen for buffer subtraction are highlighted in pale green. (C) The theoretical SAXS profile for the hP13 ZnF5-WWE1-WWE2 model from AlphaFold (in purple) is overlaid on the experimental SAXS data for hP13 ZnF5-WWE1-WWE2 shown as log(intensity) versus scattering angle (q) (in grey). (D) The probability distribution of interatomic distances, P(r), as calculated for the hP13 ZnF5-WWE1-WWE2 AlphaFold model (in purple) is overlaid on the P(r) calculated for the hP13 ZnF5-WWE1-WWE2 experimental data (in grey). (E) Dummy atom model from DAMMIF/IN analysis of mP13 scattering data, aligned with the compact monomer of mP13 ZnF5-WWE1-WWE2. (F) Dummy atom model from DAMMIF/IN analysis of hP13 scattering data, aligned with the AlphaFold hP13 ZnF5-WWE1-WWE2 model.

**Supplementary Figure 2: Sequence Alignment for PARP13 ZnF5-WWE1-WWE2.** The indicated PARP13 sequences covering the ZnF5-WWE1-WWE2 region were aligned using ESPript 3.0 (Robert and Gouet, 2014). Structural regions are indicated: ZnF5 (green box), WWE1 (purple box), WWE2 (yellow box), and Linker (blue box). The conserved ZnF5 CCCH residues that bind zinc(II) are indicated by green asterisk symbols. The highly conserved residues in WWE1 and WWE2 that contribute to the groove are indicated by purple asterisk symbols. The non-conserved WWE1 residues forming the loops of its non-functional binding cavity are highlighted by black circles. The residues in the highly conserved ends and the moderately conserved center region of the Linker connecting WWE1 to WWE2 are displayed as black and blue triangles, respectively.

## Notes

### Competing Interest Statement

The authors have declared no competing interest.

