## Supplemental Figure 1 and 2 for "Crystal structures and functional analysis of the ZnF5-WWE1-WWE2 region of PARP13/ZAP define a new mode of engaging poly(ADP-ribose)"

#### Supplementary Figure 1

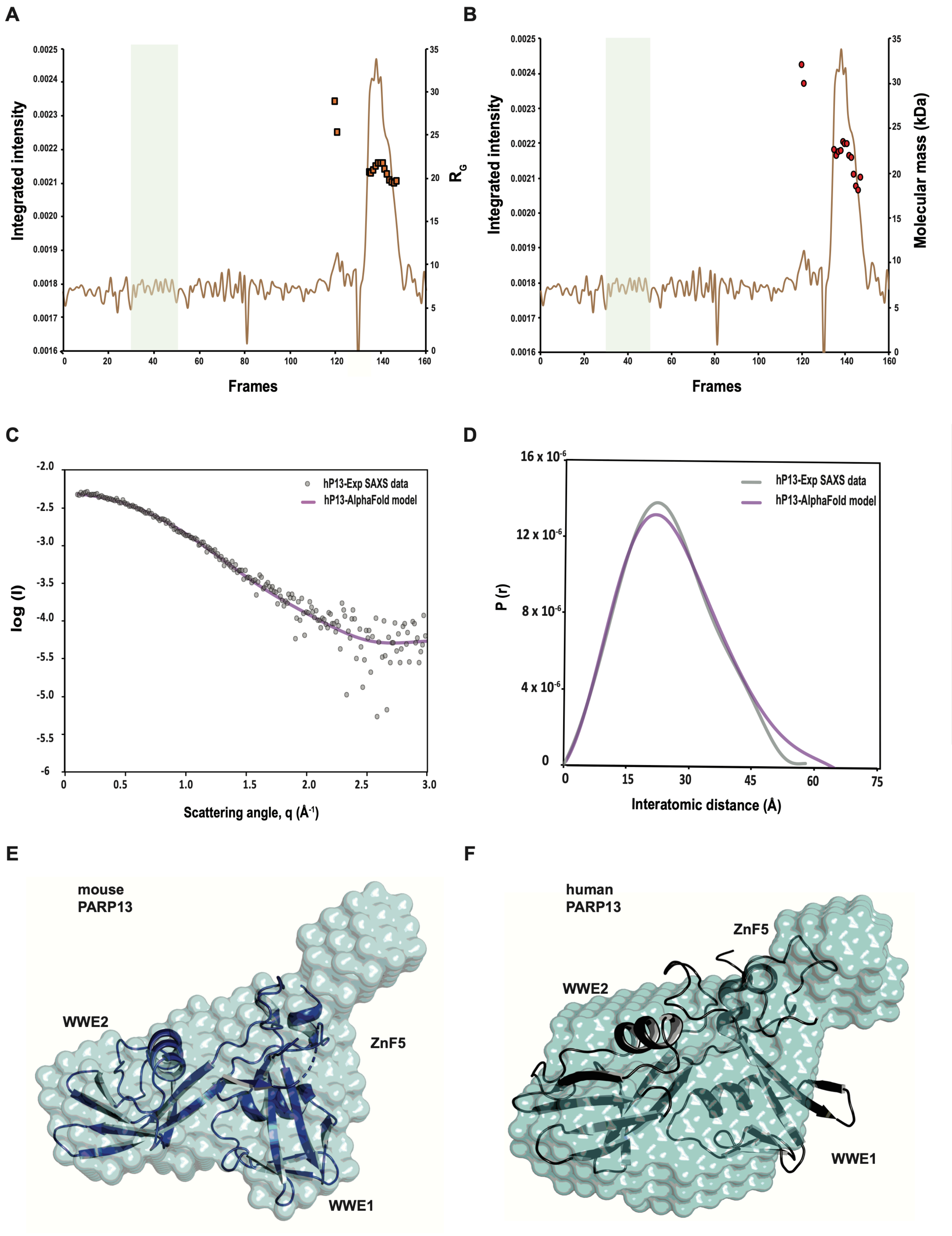

### Supplementary Figure 2

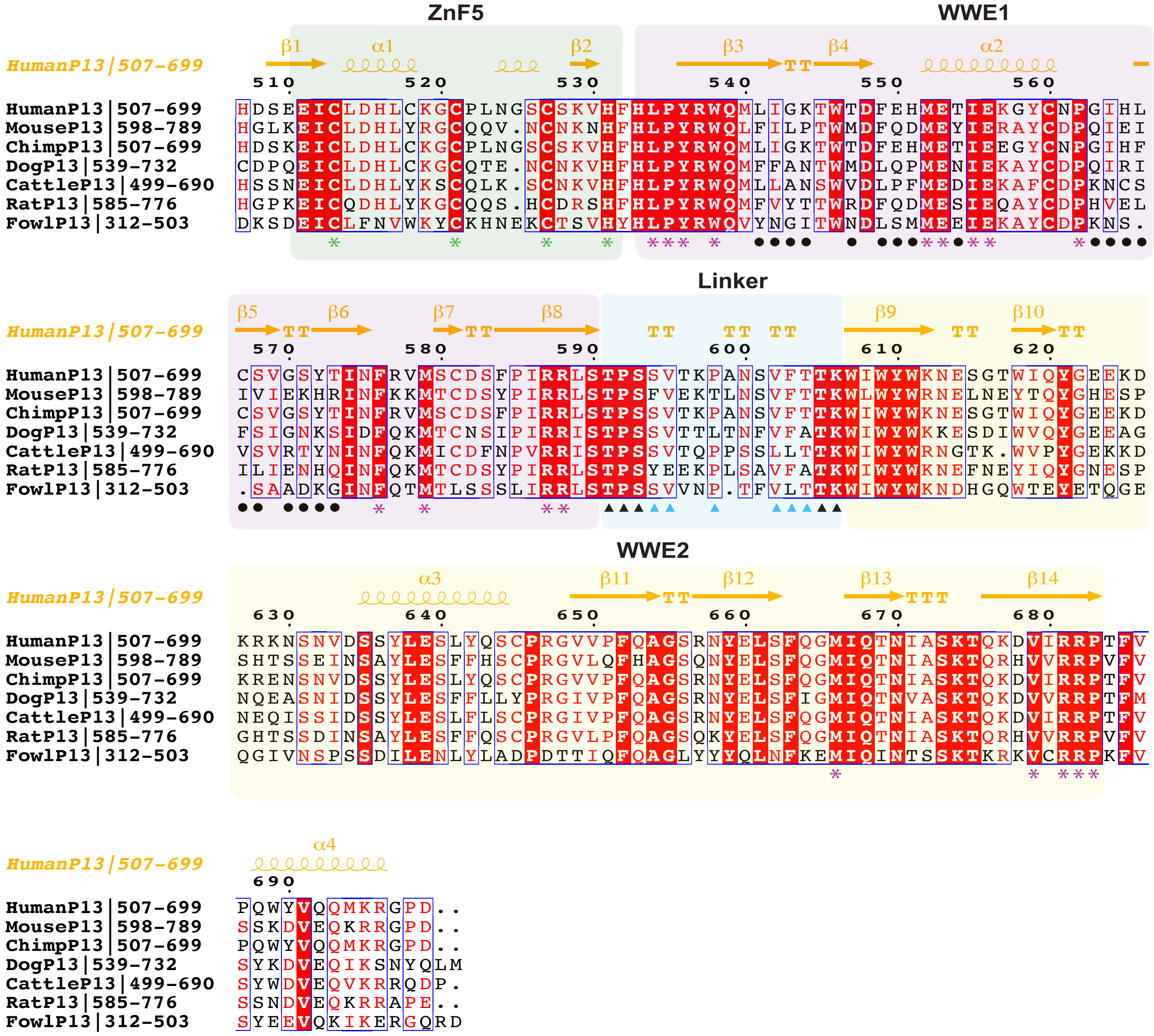
